# Rapid quantitative assessment of fish larvae community composition using metabarcoding

**DOI:** 10.1101/2020.01.01.884544

**Authors:** Frances C. Ratcliffe, Tamsyn M. Uren Webster, Deiene Rodriguez-Barreto, Richard O’Rorke, Carlos Garcia de Leaniz, Sofia Consuegra

**Affiliations:** Department of Biosciences, College of Science, Swansea University, Swansea SA2 8PP, United Kingdom

**Keywords:** Bulk samples, correction factors, fish larvae, metabarcoding, quantitative

## Abstract

Climate change stressors greatly impact the early life-stages of many organisms but their cryptic morphology often renders them difficult to monitor using morphological identification. High-throughput sequencing of DNA amplicons (metabarcoding) is potentially a rapid and cost-effective method to monitor early life-stages for management and environmental impact assessment purposes. Yet, there is conflicting information about the quantitative capability of metabarcoding. We compared metabarcoding with traditional morphological identification to evaluate taxonomic precision and reliability of abundance estimates, using 332 fish larvae from multinet hauls (0-50m depth) collected at 14 offshore sampling sites in the Irish and Celtic seas. To improve relative abundance estimates, the amount of tissue for each specimen was standardised and mitochondrial primers with conserved binding sites were used. Family level correction factors for amplification bias and back-calculations were applied to estimate numbers of individuals of a given taxon in a sample. Estimates from metabarcoding reads and morphological assessment were positively correlated for relative family abundances as well as taxon richness (Rs=0.81, P=0.007) and diversity (Rs=0.88, P=0.003). After applying family level correction, back-estimates of the number of individuals per family within a sample were accurate to ± 2 individuals. Spatial patterns of community composition did not differ significantly between metabarcoding and morphological assessments.

Our results show that DNA metabarcoding of bulk tissue samples can be used to monitor changes in fish larvae abundance and community composition. This represents a feasible, efficient and faster alternative to morphological identification that can be applied to terrestrial and aquatic habitats.

## Introduction

Over the course of this century, climate change is expected to become the greatest driver of change in both terrestrial and marine biodiversity (Garciá Molinos et al., 2016). The sensitivity of organisms’ early life stages to abiotic stressors plays a central role in driving ecological responses to climate change (Radchuk, Turlure, & Schtickzelle, 2013). Therefore, monitoring of larval organisms provides critical information to understand these responses over time (Asch, 2015), informing conservation measures, policy and regulation (Ellis, Milligan, Readdy, Taylor, & Brown, 2012; Borja, Elliott, Uyarra, Carstensen, & Mea, 2017). However, the cryptic morphology of early life-stage organisms can render them difficult to monitor using morphological identification (Sigut et al., 2017), as, for example, in the case of fish larvae (Brechon, Coombs, Sims, & Griffiths, 2013; Kimmerling et al., 2018).

Traditional fish larvae monitoring involves identifying each individual using a light microscope, counting myotomes, assessing pigmentation patterns and jaw morphology (Russel 1976). Yet, identification keys are incomplete for many parts of the world (Becker, Sales, Santos, Santos, & Carvalho, 2015) and, where descriptions are available, morphological assessment is time consuming and requires specialist training (Brechon *et al*., 2013). Morphological taxonomy also relies on the identifying features remaining intact for species level assignment (Russel, 1976), but damage is common during sampling (e.g. continuous plankton recorders), leading to misidentification and loss of valuable information (Richardson et al., 2006).

In cases where morphological identification is unfeasible, DNA sequencing technologies may be used to identify organisms (Taberlet, Coissac, Pompanon, Brochmann, & Willerslev, 2012). The development of high-throughput sequencing technology allows amplicon-based sequencing (metabarcoding) of multiple individuals of various species concurrently (i.e. bulk samples), providing a relatively quick method of processing many samples to obtain taxonomical information (Taberlet et al., 2012). However, obtaining accurate abundance estimates through relative read abundance (RRA) from amplicon sequence data has remained challenging (Deagle et al., 2019; Lamb et al., 2019). This is because biases in RRA estimations can be introduced at different stages of the metabarcoding protocol, for example, cell and DNA quantity, extraction success and PCR amplification rates can vary between tissue type and species (Lamb et al., 2019), leading to inaccurate estimates. Another source of bias can arise from unequal body size of individuals pooled within a bulk sample, which can be mitigated by size fractionation of organisms prior to extraction, increasing the reliability of RRA estimates (Elbrecht et al., 2017). The choice primers and target region may introduce further bias (Deagle, Jarman, Coissac, Pompanon, & Taberlet, 2014). These biases have led to designing costly and bioinformatically challenging metagenomic approaches (Tang et al., 2015; Kimmerling et al., 2018) or to the use of multiple loci (Richardson et al., 2015) to estimate abundance.

Improving the reliability of abundance estimates is thus needed to make metabarcoding more useful for biodiversity monitoring, calculation of metrics such as diversity indices, as well as detection of shifts in multispecies community composition (Bohmann et al., 2014). Different approaches have been proposed to improve abundance estimates based in RRA, whilst still using a cost effective, single marker PCR approach (Thomas, Deagle, Eveson, Harsch, & Trites, 2016; Elbrecht, Peinert, & Leese, 2017). For example, using primers with widely conserved priming sites may reduce taxa specific biases (Krehenwinkel et al., 2017), although taxonomic resolution can be reduced due to highly similar sequences within a family (Thomsen, Møller, et al., 2016). Methods such as taxa specific tissue correction factors (Thomas, Jarman, Haman, Trites, & Deagle, 2014; Thomas et al., 2016b) may also be used to back-estimate relative proportions of tissue from RRA and with that estimate numbers of individuals in a sample.

Here, using a single mitochondrial marker (12S ribosomal RNA), we improved the reliability of DNA metabarcoding abundance estimates by standardizing input material, choosing conserved primer binding sites and applying correction factors for amplification bias. Using bulk fish larvae samples from the Irish and Celtic Seas, we compared the sensitivity and accuracy of this approach with traditional morphological identification, to assess whether metabarcoding can be a feasible and rapid alternative to traditional assessment for estimating fish larvae richness, diversity and community composition metrics.

## Materials and Methods

### Field sampling

Sampling was carried out onboard the RV Celtic Voyager between May 17^th^ and May 26^th^ 2018. Fish larvae (3-30mm) from 14 hauls (1 oblique haul to 50m depth per site) were sampled using a MultiNet plankton sampler (Hydro-Bios, Kiel, Germany), filtering a mean volume of 215 ± 55 m^3^ of water for hauls 1-8 and 12. Hauls 9-14 (with the exception of haul 12) were sampled in two vertical hauls, from the surface to 50m, filtering a mean volume of 38 ± 6m^3^, both hauls were pooled. Fish larvae from each haul were separated from other zooplankton species and preserved in RNAlater at room temperature for 24hrs, then refrigerated at 4°C until morphological identification.

### Morphological Identification

Fish larvae ranged from 2mm-30mm total length. For morphological identification larvae were first separated into major groupings following Russel (1976) and subsequently assigned to family level. Assignment to genus, and species where possible, was then carried out. Assignments were checked against the species descriptions first in Russel (1976), and, where possible, double checked against the description by Rodriguez et al. (2017). If the taxa of a particular individual could not be confidently assigned, the specimen was pooled with individuals of similar morphology and a specimen of the group added to a reference collection. The reference collection contained 32 individuals subsampled across all 9 hauls and was used test morphological identification accuracy and correct morphological taxonomic assignments. The collection was barcoded using Sanger sequencing using the 12S V5 primers (Riaz et al., 2011), and taxonomy was assigned using the MegaBLAST algorithm (Morgulis et al., 2008) against the National Center for Biotechnology Information (NCBI) GenBank nucleotide database (accessed November, 2018). Where species level identification could not be resolved with 12S V5, due database limitations or synonymous sequences, CO1 barcoding (Ward, Zemlak, Innes, Last, & Hebert, 2005) was also used. The reference collection was used to calculate the accuracy of taxonomic identification to family, genus and species levels. In cases when the morphological and barcoding assignments differed in the reference collection, barcoding identities were used to correct the morphological assignments for 75 of the 332 larvae for downstream analysis, referred to as Sanger corrected identities. Ammodytidae and Clupeidae were not discriminated using this method, due to similar morphologies preventing accurate groupings within these families. Additionally, a separate sub-group of specimens (15) were analysed by a second taxonomist to estimate accuracy and repeatability.

### DNA extraction

After taxonomic identification, bulk tissue samples for each haul were prepared for DNA extraction as follows: 5mg (+/-3mg) of tissue were cut from the area anterior to the tail of each larva (for individuals <5mg, the entire larva was used) and placed in a falcon tube on ice. Buffer ATL and proteinase K (Qiagen DNeasy Blood and Tissue kit) were then added to the pooled tissue sample in a ratio of 180μl of ATL and 20μl proteinase K for 15mg of tissue. Each falcon tube (representing a haul) was vortexed thoroughly and incubated overnight to digest at 56°C, shaking at 65 rpm. Samples were visually inspected for tissue remnants, vortexed and re-incubated until all tissue dissolved. Digestions from each haul were then vortexed for 45 seconds to ensure thorough mixing of digested products and divided in three sub-samples of 200ul that were extracted using the Qiagen DNeasy Blood and Tissue kit, following the manufacturer’s instructions. Extraction blanks were carried through each step of the process.

To ascertain whether taxa were subject to PCR bias, two mock communities were generated by pooling known concentrations of genomic DNA (quantified using Qubit dsDNA HS Assay (Invitrogen) from the taxa identified using Sanger sequencing. Community 1 contained equimolar concentrations of each of the following taxa: *Ammodytes marinus, Callionymus sp, Ciliata mustela, Sprattus sprattus, Lepidorhombus sp, Merluccius merluccius, Merlangus merlangius, Tricopterus minutus*. Community 2 contained *A. marinus* (1.83%), *Callionymus sp* (23.77%), *C. mustela* (2.64%), *C. harengus* (1.66%), *Lepidorhombus sp* (18.78%), *M. merluccius* (22.10%), *M. merlangius* (26.92%), *T. minutus* (2.30%). These samples were subjected to the library preparation, sequencing and analysis workflow alongside the bulk samples.

### Library preparation and sequencing

A 106 bp fragment of the 12S mitochondrial was amplified with the 12S V5 primers (Riaz et al., 2011) using Phusion^®^ High-Fidelity DNA Polymerase (New England Biolabs Ipswich, MA, USA), with an annealing temperature of 52°C, in 3 extraction replicates per haul. Libraries were prepared using a 2-step PCR approach, based on the Illumina 16S Metagenomic Sequencing Library preparation guidelines (Illumina, Inc., San Diego, CA, USA), with following adaptations: in the first PCR step, each extraction replicate was triplicated in order to increase detection of rare species (Alberdi, Aizpurua, Gilbert, & Bohmann, 2018). Subsequently, 10ul from each triplicate were pooled prior to first cleanup. Cleanups were performed using Agencourt AMPure XP beads (Beckman Coulter, Brea, CA, USA), using a 1.2 x volume of beads to PCR product. Amplicons were indexed using Nextera XT Index Kit v2 Set C (Illumina, Inc., San Diego, CA, USA), and DNA concentration of each reaction was quantified via Qubit dsDNA HS Assay (Invitrogen, Carlsbad, CA USA) and pooled in equal molar concentrations. PCR and extraction blanks (using molecular grade water instead of template) were subjected to all steps of the library preparation process. In addition, a sequencing/tag jumping blank, where no sample was added prior to sequencing, was used. Pair-end sequencing was carried out using the Illumina MiSeq platform (Illumina, San Diego, CA, USA).

### Bioinformatics-sequence processing

De-multiplexed samples containing raw pair end sequences were processed using Qiime2 (version 2019.1, Bolyen et al., 2019). Initially, raw sequences were quality checked using interactive quality plots, in order to obtain values for sequence trimming and truncation. De-noising was carried out using DADA2 (Callahan, McMurdie, & Holmes, 2017) where the first 10 bp of each sequence were trimmed to remove adaptors and all sequences truncated to 100 bp in length based on quality scores. Default DADA2 settings within Qiime2, were used to detect and, where possible, correct sequencing errors and filter out phiX reads, and chimeric sequences, join pair end reads and de-replicated sequences. The amplicon sequence variant (ASV) approach was chosen because it provides a higher resolution than a traditional OTU approach, enabling detection of single nucleotide differences (Callahan et al., 2017). After de-noising, the ASV and BIOME tables were exported for taxonomic assignment.

### Database construction and taxonomic assignment

A custom database was constructed using *in silico* PCR against the NCBI database (downloaded February 2019): primers were allowed to have 3 base mismatches *in silico* (search_PCR command, Edgar, 2010) and a corresponding taxonomy file was constructed using the obiannotate tool (OBITools, Boyer et al., 2016). All sequences were trimmed to the target region. A list of all marine fish species encountered in the British Isles, including non-native fish (366 species) (Fish Base: accessed 31/3/2019) was then used to filter the main database to fish species present in the study region, of which 207 were available. Additionally, 6 Sanger sequences from the reference collection in this study were added. The database included marine mammals, bacteria and other contaminants that might be amplified by the primers such as *Homo sapiens*.

Initially, ASVs were classified using the KNN method in Mothur (Schloss et al., 2009) using parameter ‘numwanted=1’ (Findley et al., 2013), against the custom database. Because this parameter may lead to false positive assignments, KNN assignments were then verified using NCBI megaBLAST, with max-target sequences =10. The top 10 assignments were screened for UK species (Fish Base) on a case by case basis. Where the percentage of UK species match fell below 98%, or where multiple UK species matched above a 98% match, MEGAN (6.15.1) was were used to assign species to the lowest common ancestor (Huson, Auch, Qi, & Schuster, 2007). ASVs for which there were no vertebrate matches were discarded from downstream analysis. A subsample of the assignments (32 specimens) were then checked, using the sequences from the reference collection, to ascertain if they were present in the correct haul and taxonomy was correctly assigned (100% were correct).

Tag jumping/cross contamination (Schnell, Bohmann, & Gilbert, 2015) was removed on the following basis: a taxa was removed from a haul if it had less than 115 reads (maximum reads for a single species in tag jumping control sample) or did not appear in all 3 replicates.

Family correction factors (FCFs) for sequencing bias were calculated based on (Thomas et al., 2014) for each family in each haul, using the following formula:

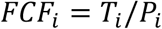

Where *i* is the family of interest, *T_i_* is the proportion of the family in the bulk tissue sample, and *P_i_* is the proportion in the haul amplicon pool. FCFs were calculated for a given family in a given haul and then averaged to provide a mean correction factor for each family (mFCF). For spatial analysis, numbers of individuals of taxon in a haul were back-estimated from the proportion of reads in the corresponding sample, as follows:

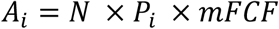

Where *A_i_* is the abundance (number of individuals) of the taxon of interest (*i*) in a given haul, *N* is total number of individuals in the haul, *P_i_* is the proportion of that taxon in the haul amplicon pool and *mFCF* is the mean tissue correction factor for the corresponding family.

### Statistical analysis

The accuracy of estimates of RRA and diversity indices derived from metabarcoding was assessed against results from morphological taxonomy using Spearman’s rank correlation analysis performed in R (version 3.5.2). All diversity index calculations were performed on RRA and morphological relative abundances. Mock community abundances (quantity of DNA inputted and proportion of reads) were compared using Chi-Square tests. For spatial analysis, the survey area was divided into 3 locations along a temperature gradient: Loc 1 (Above the Celtic/Irish sea front, 9-10.99°C), Loc 2 (Channel spawning grounds, 11-12.99°C), Loc 3 (Western Celtic Sea, 13-14°C) (Figure 1). The number of individuals and back-estimated reads of a given taxon were divided by the volume of water filtered in the corresponding haul (Canfield & Jones, 1996) to obtain catch per unit filtered (CPUF) or back-estimated abundances from reads per unit filtered (RPUF) values respectively. The family Ammodytidae was excluded from this analysis, because not all individuals were retained in haul 4. CPUF and RPUF values were square-root transformed, and composition similarity calculated by hierarchical clustering using a Brae-Curtis resemblance matrix. Subsequently, pairwise analysis of similarities (ANOSIM) were used to test whether there was a significant difference in community composition between locations (Clarke, 1993), using both the CPUF and RPUF methods. Where significant differences were detected, SIMPER analysis (Clarke, 1993) was used to ascertain which taxa accounted for the differences observed. Diversity indices calculations and multivariate spatial analyses were performed using Primer-v7 (Clarke & Gorley, 2015).

**Figure 1.**
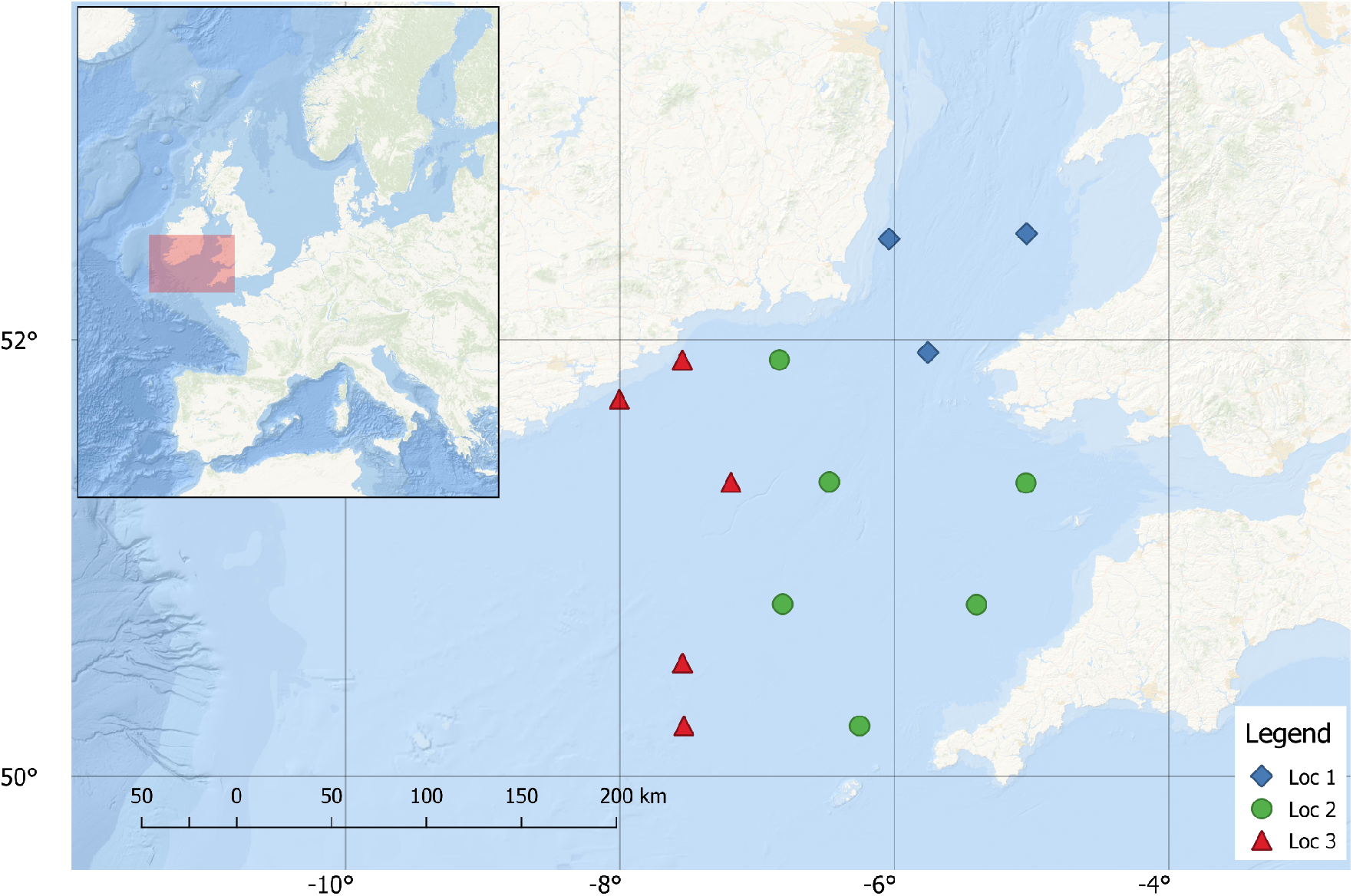
Mulitnet haul locations in the Irish and Celtic seas. Locations for spatial analysis, based on SST, are indicated as Loc 1 (Above the Celtic/Irish sea front: 9-10.99°C), Loc 2 (Channel spawning grounds: 11-12.99°C), Loc 3 (Western Celtic Sea: 13-14°C).

## Results

A total of 332 fish larvae were caught in 11 of the 14 hauls in the survey. No larvae were encountered in hauls 10, 11, and 14 and only one in hauls 1 and 6, therefore 9 of the 14 hauls were used in metabarcoding. The maximum number of individuals per haul was 63 (haul 2) (Table S1).

Morphological identification assigned 325 (98%) of individuals to family level. It was not possible to assign the families of the remaining 8 larvae, due to damaged identifying features. Of those specimens assigned to family level, 255 (77%) were assigned to a genus and 100 (30%) to a species. The reference collection (comprising of 32 individuals across 9 hauls), contained the following species: *A. marinus, S. sprattus, T. minutus, Limanda limanda, C. mustela, Clupea harengus, Trisopterus esmarkii, Labrus bergylta, Callionymus sp, Microstomus kitt, Buenia jeffreysii, Lepidorhombus sp., M. merluccius and M. merlangus*. Of these individuals, morphological identification correctly assigned 76.47% of the specimens to family, 32.35% to genus and 17.65% to species. For the 15 individuals morphologically identified by two independent observers before Sanger sequencing, they were 100% and 93% correctly assigned to family level by the first and second observer respectively, 86.3% and 53.3% to genus and 40% to species level in both cases.

A total of 3,398,391 raw pair end reads were generated for this study. After Qiime2 DADA denoising, a total of 2,675,140 reads remained for downstream analysis. Once the taxonomic assignment was complete, reads likely present due to tag jumping from concurrent sample sequencing (*Solidae* 274 reads, *Scomber scombrus*, 10 reads, *Salmo salar*, 3 reads), and human reads (2338) were removed from downstream analysis. A total of 49 fish ASVs remained downstream analysis. Samples contained a mean of 93,223 reads (standard deviation = 31,866) post filtering, the tag jumping blank contained 146 reads, the PCR blank 64 reads and extraction blanks 116 and 71 respectively. Tag jumping read removal resulted in 0.046% of reads being excluded from downstream analysis across the samples in the study. Post filtering, taxa distribution was concordant among the 3 haul replicates in all 9 hauls (Figure 3).

Sequencing of the mock communities 1 and 2 resulted in 90,199 and 58,690 reads respectively. All 8 species added to the mock communities were detected using metabarcoding and only reads pertaining to the input DNA were observed, with the exception of 23 reads assigned to *T. esmarkii* in Mock community 1. Mock community 1, for which equimolar concentrations of DNA had been pooled, contained a mean number of 11,272 reads per species (standard deviation: 4,603 reads) and the relative quantity of DNA inputted differed from the RRA expected (Chi-square: X^2^ = 14.59, df = 7, p-value = 0.04). For Mock community 2, where input molar concentrations were varied, the relative input did not differ from the RRA (X^2^ = 11.39, df = 7, p-value = 0.12, Figure S1).

### Comparison of taxonomy and abundance estimates by morphology and metabarcoding

The relative abundance of individuals identified morphologically in a sample and the corresponding RRAs were positively correlated for all families assessed (Ammodytidae R_s_ =0.93, P<0.001, Callionymidae R_s_ = 0.99, P<0.001, Clupeidae R_s_ = 0.97, P<0.001, Gadidae R_s_ = P<0.001, Pleuronectidae R_s_ = 0.68, P = 0.05, Triglidae R_s_ = 0.88, P = 0.002, Figure S2). In addition, no difference in diversity and taxon richness were detected between the relative abundance of morphological assignments and RRA assignments (Spearman’s Rank: richness: Rs = 0.81, P=0.007, Shannon Index: Rs 0.88, P=0.003, Simpson’s Diversity: Rs = 0.8, P= 0.01, Figure 2).

**Figure 2.**
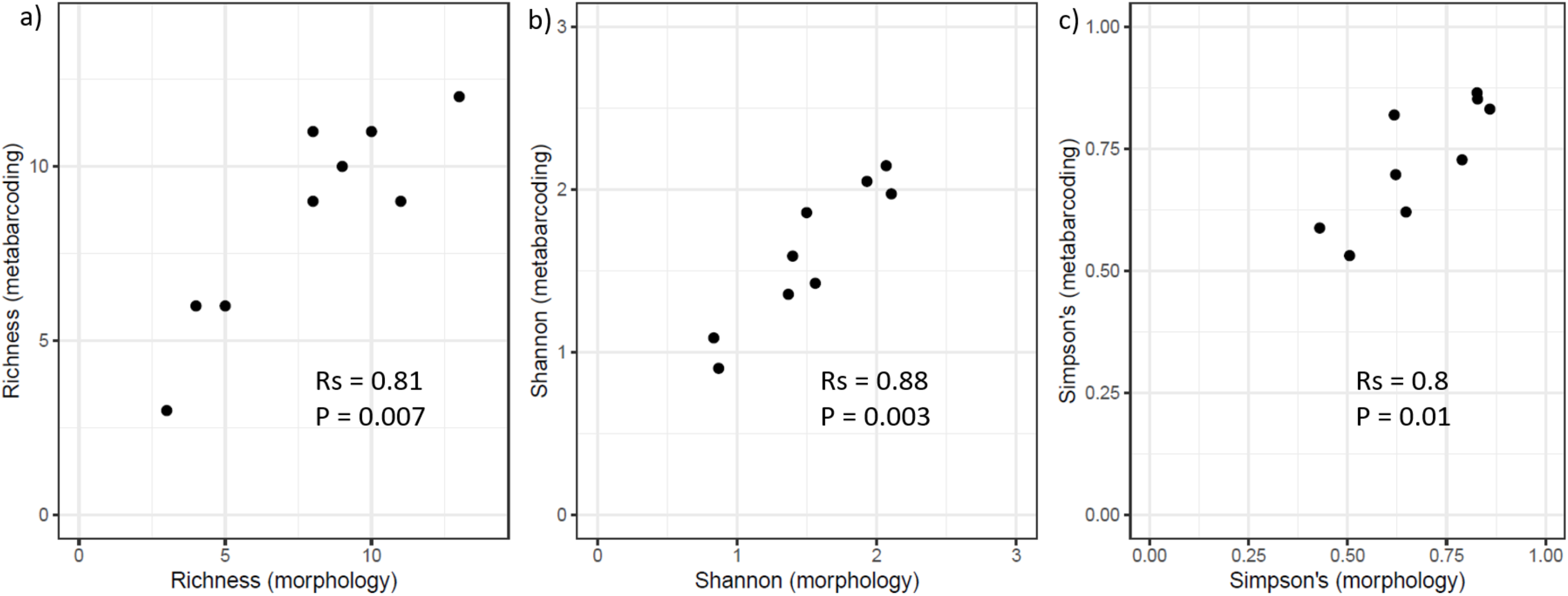
Consistency of diversity metrics between relative abundances of morphological assignments and relative read abundance assignments, post bioinformatic filtering, for a) species richness, b) Shannon wiener diversity index and c) Simpson’s diversity (1-lamda). Rs = Spearman’s rank Rho values.

**Figure 3.**
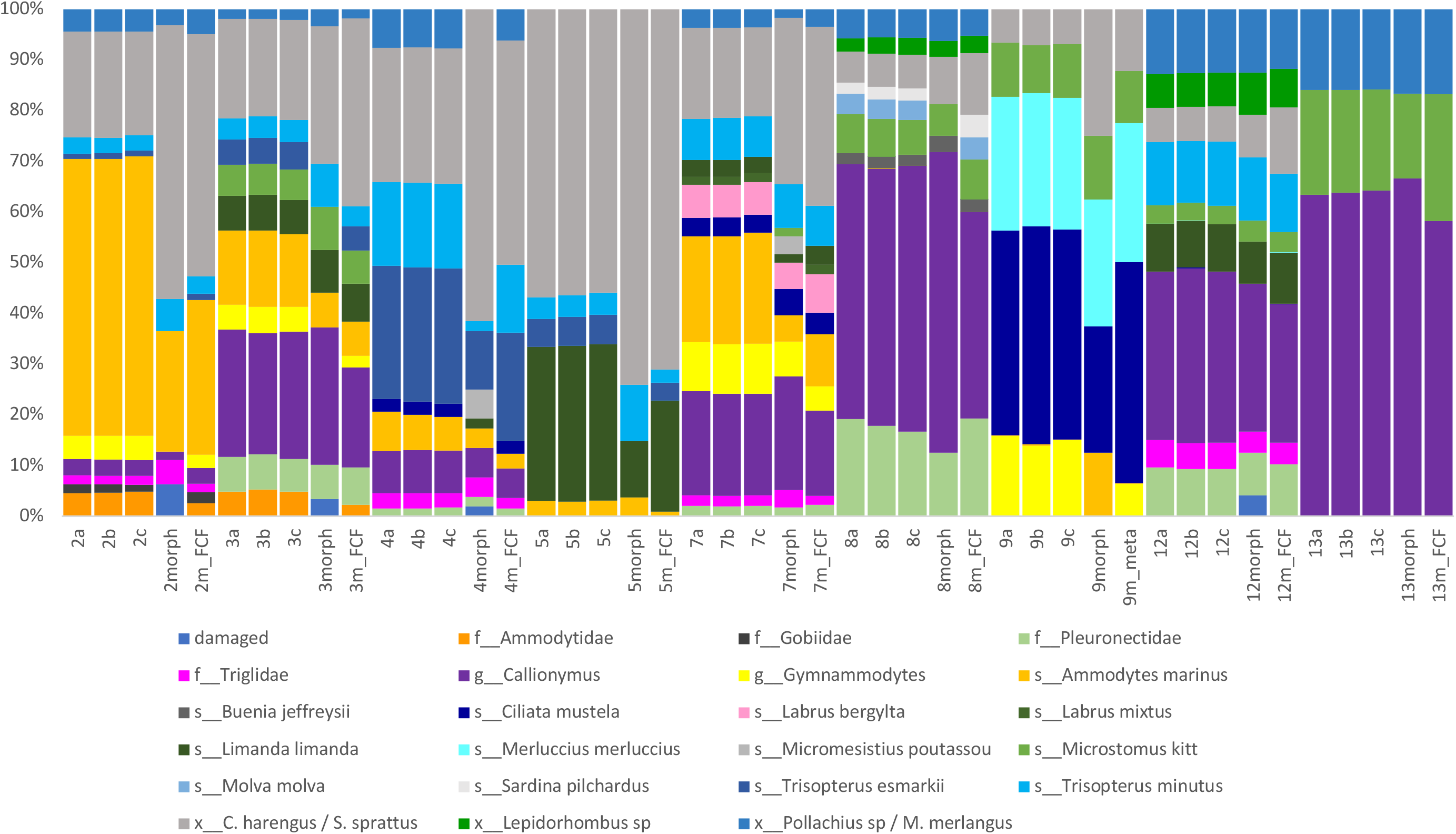
Comparison of relative read abundances before back-estimation, (3 replicates per haul, ‘a’, ‘b’, ‘c’ samples) and morphological taxonomic assignments, corrected by Sanger sequencing (1 per haul, ‘morph samples’). f__indicates family level assignment, s__indicates species level and x__indicates 2-3 possible species assignments. Morphological assignments of *P. pollachius/viens, M. merlangus* were grouped and morphologically assigned *Glyptocephaus cycnoglossus* has been re-assigned to Pleuronectidae to match metabarcoding assignments to aid visual interpretation of abundances.

Based on morphology alone, before correction by Sanger barcoding, taxa within Ammodytidae and Clupeidae could not be assigned further than family level. Incorrect morphological assignments occurred in the cases of *Micromesistius poutassou* (Sanger seq: *M. merlangius*), Aphia minuta (Sanger seq: *C. harengus/ S. sprattus*) and *Mugilidae* (Sanger seq: *L. bergylta* and *C. mustela*) (Figure 4). In addition, *Sardina pilchardus, Labrus mixtus/bergylta, Molva molva*, and a taxon belonging to the Gobiidae family, could only be detected using sequencing. In contrast, *M. merlangus*, and *Pollachius virens/pollachius*, were identifiable through morphology, but not resolved to species level by metabarcoding due to lack of variability of the 12S fragment. *C. harengus* and *S. sprattus* could not be separated by morphology or metabarcoding.

**Figure 4.**
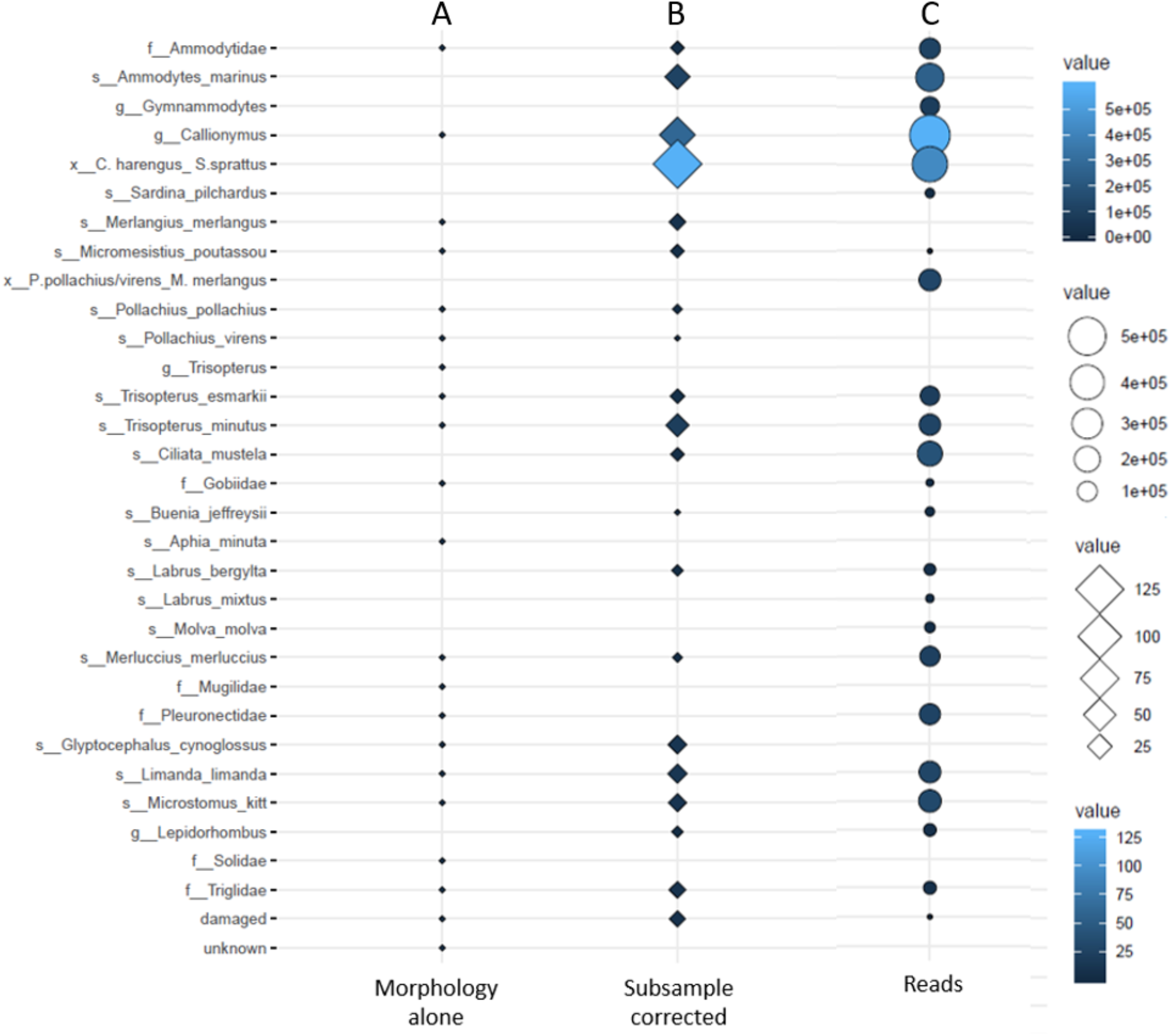
Overview of larval detections during the survey. Panel A: Taxonomic assignments using morphology alone (presence/absence). Panel B: Morphological taxonomic assignments corrected using a Sanger sequenced subsample (reference collection), diamonds represent total number of larvae of a taxa observed during the survey. Panel C: Metabarcoding taxonomic assignments, circles represent total number of reads obtained for each taxa, post-filtering.

We calculated mFCFs to further improve abundance estimates by comparing the proportions of the 6 most abundant families in the tissue mix to the proportions observed in the amplicon pool on a haul by haul basis (Table 1). Ammodytidae was the most strongly overrepresented, resulting in the smallest mFCF (0.41) and Clupeidae was the most underrepresented, resulting in the largest (1.69). When these mFCFs and back-estimation were applied to the RRA in a given haul, the mean difference between the number of morphologically assessed Ammodydidae and the number of individuals back-estimated from RRA was reduced to −0.54 (SD: 1.58) individuals, a mean improvement of 8.38 individuals per haul. Clupeidae abundance back-estimates improved by mean 5.38 individuals, resulting in a mean difference of 1.04 (SD: 5.48) individuals per haul. Improvements were less marked for the remaining families (Table 1). For families which were not strongly under/over represented (Triglidae and Callionymidae), mFCFs did not improve back-estimates and were therefore not applied in RPUF calculations.

**Table 1.**
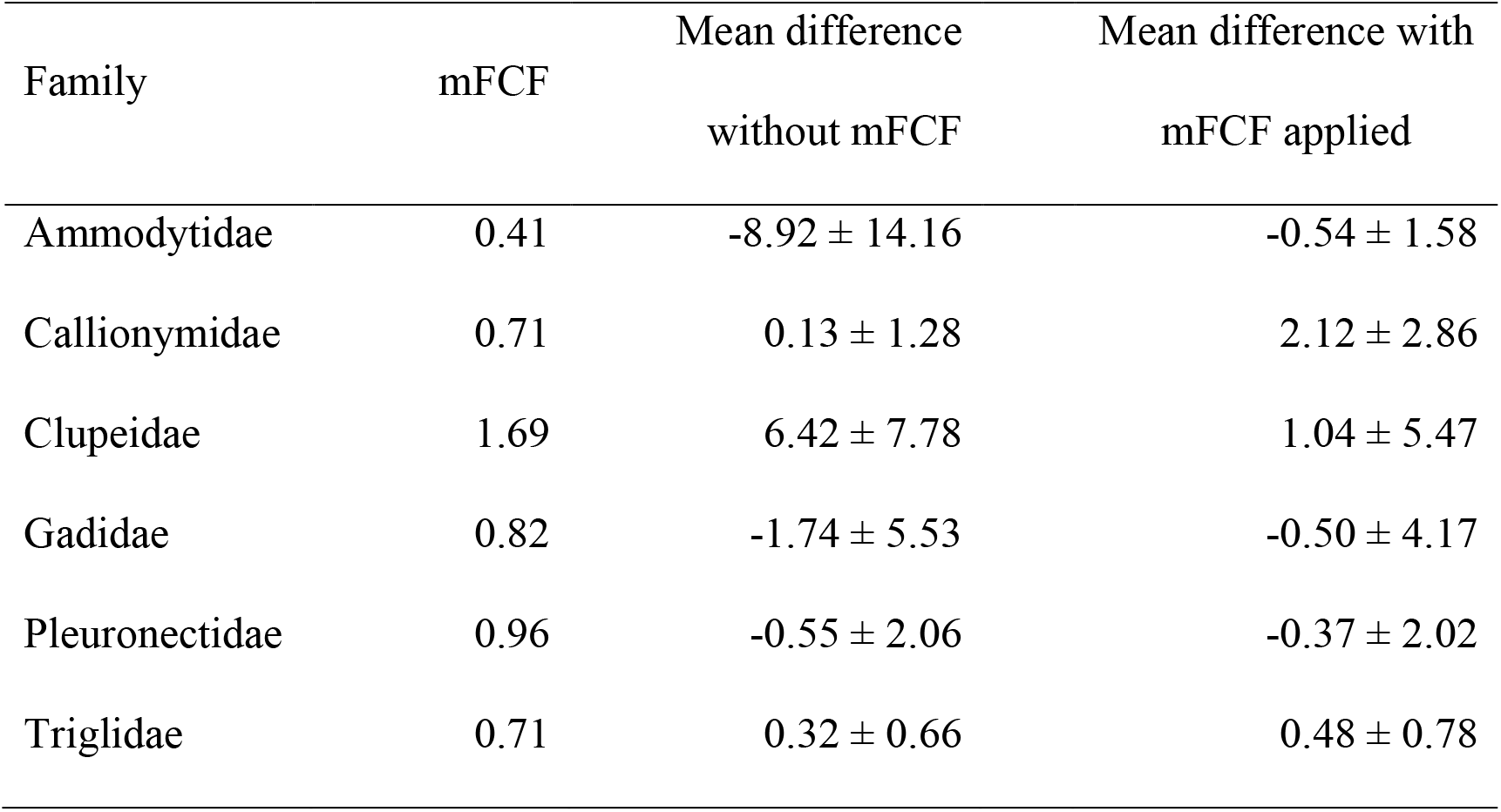
Mean Family level Correction Factors (mFCFs) for 6 families. Mean differences in abundance (number of individuals) between morphology and back-estimated reads across all samples in the study are represented with and without correction factor applied, ± standard deviation.

| Family | mFCF | Mean difference<br>without mFCF | Mean difference with<br>mFCF applied |
| --- | --- | --- | --- |
| Ammodytidae | 0.41 | $-8.92 \pm 14.16$ | $-0.54 \pm 1.58$ |
| Callionymidae | 0.71 | $0.13 \pm 1.28$ | $2.12 \pm 2.86$ |
| Clupeidae | 1.69 | $6.42 \pm 7.78$ | $1.04 \pm 5.47$ |
| Gadidae | 0.82 | $-1.74 \pm 5.53$ | $-0.50 \pm 4.17$ |
| Pleuronectidae | 0.96 | $-0.55 \pm 2.06$ | $-0.37 \pm 2.02$ |
| Triglidae | 0.71 | $0.32 \pm 0.66$ | $0.48 \pm 0.78$ |

### Spatial distribution of larvae assessed by both methods

Catch per unit filtered (CPUF) and back-estimated reads per unit filtered (RPUF), with mFCFs applied to Clupeidae, Gadidae and Pleuronectidae families, were no different between locations 1 and 2, and 1 and 3, although locations 2 and 3 differed in composition (ANOSIM CPUF R=0.233, P = 0.039, RPUF R=0.218, P = 0.037, Table 2). SIMPER analysis (% dissimilarity contribution) attributed 48.35% (CPUF) and 44.87% (RPUF) of the difference in composition between locations 2 and 3 to three taxa: *C. harengus/S. sprattus* (CPUF: 21.44%, RPUF: 19.91%), *Triglidae* (CPUF: 14.37%, RPUF: 14.05%) and *Callionymidae* (CPUF: 12.54%, RPUF: 10.91%) (Table S3). The greatest difference observed in dissimilarity contributions for the remaining, less abundant taxa was 2.7% (*C. mustela*).

**Table 2.** ANOSIM matrix, showing R values of pairwise comparisons of community composition between 3 locations in the Irish/Celtic seas, using morphological taxonomic assignments and abundances (CPUF sanger corrected) and metabarcoding taxonomic assignments and back-estimated abundances (RPUF). * indicates significant difference in community composition between 2 locations.

|  |  | RPUF |  |  |
| --- | --- | --- | --- | --- |
|  |  | Loc 1 | Loc 2 | Loc 3 |
| CPUF | Loc 1 |  | -0.105 | -0.027 |
|  | Loc 2 | -0.111 |  | 0.218* |
|  | Loc 3 | -0.053 | 0.233* |  |

## Discussion

Here we demonstrate that metabarcoding is a reliable and practical alternative to traditional morphological assessment. We show that accurate RRA estimates can be achieved by standardising the amount of tissue analysed per specimen, choosing primers with conserved binding sites and using family level correction factors. These estimates can then be used to successfully calculate numbers of individuals and community composition metrics needed to monitor changes over time.

There is considerable debate over whether amplicon sequencing can deliver reliable quantitative data (Deagle et al., 2019). The reliability of abundance estimates from metabarcoding varies considerably between studies, with some showing only a weak correlation between RRA and abundance (Lamb et al., 2019; Piñol, Senar, & Symondson, 2019). However, useful information may be obtained from the RRA within a sample, rendering this approach often more informative than presence/absence assessment (Deagle et al., 2019). In contrast, metagenomic approaches that do not require PCR amplification can successfully estimate abundance (e.g. Kimmerling et al., 2018), although the costs and bioinformatic complexity of this approach may be prohibitive in many contexts (Porter & Hajibabaei, 2018). Therefore, improvement of metabarcoding abundance estimates enables wider application of amplicon sequencing.

We have shown that the use of approximately equal weights of tissue per individual can improve RRA and diversity estimates. By using small pieces (~5mg) of tissue, we were also able to eliminate the grinding/cryogenic grinding step when preparing samples for extraction, saving time and reducing the risk of contamination between samples which may occur when using grinding tools. Approaches based on photographically assessing the surface area of taxa and modelling biomass might also eliminate the need for weighing tissue (Kimmerling et al., 2018), although not necessarily reducing time and costs.

While is no perfect marker for all studies (Deagle et al., 2014), we have shown here the benefits of using primers with well conserved binding sites, particularly for RRA estimates. Whereas the CO1 marker has extensive sequence databases as well as a strong capability to discriminate between species, it also carries an increased risk of amplification bias due to the lack of conserved binding sites across a broad range of taxa (Deagle et al., 2014). This can result in false negatives were taxa known to be present in a sample do not amplify (Collins et al., 2019; Nobile et al., 2019). Using more conserved priming sites, such as the 12S marker, may reduce taxa specific biases (Krehenwinkel et al., 2017), although it has been argued that taxonomic resolution may be reduced due to lack of sequence variability within families (Thomsen et al., 2016), and the completeness of reference databases also influences the resolution to species level (Miya et al., 2015).

In comparison to morphological identification without the assistance of Sanger sequencing, metabarcoding achieved higher taxonomic resolution and more accurate identifications to family level. However, morphologically assessed groupings supported by barcoding using Sanger reference sequencing achieved a similar level of assignment at the family level to metabarcoding across the study. Yet, while short reads can struggle to resolve some families to species level (Thomsen et al., 2016), hindering species level data interpretation, we found that the use of metabarcoding improved taxonomic assignment overall. Morphology performed better than sequencing only in the case of *Glyptocephalus cynoglossus*, due to distinct morphological characteristics, and in a few cases caused to lack of information or sequence variation at the 12S region. In general, synonymous sequences at the target region resulted in just two (e.g. *C. harengus/S. sprattus*) or three species (*M. merlangus/P. pollachius/P. virens*) not being distinguished from each other. For studies requiring species level identification taxa affected by lack of marker sequence information or variability, a qPCR approach (Brechon et al., 2013), or a family specific, multi primer approach (Riaz et al., 2011) could be easily used to refine metabarcoding assignments.

For some applications, genus level analysis provides similar diversity and community composition information than species level and would be appropriate, for example to detect responses to environmental change (Hernandez, Carassou, Graham, & Powers, 2013). In some other cases, family level analysis has been deemed sufficient to detect broadscale changes, e.g. after major environmental disturbance (Hernandez et al., 2013). Therefore, dependent on hypothesis, a single 12S marker may be sufficient.

The application of correction factors (Thomas et al., 2014; Thomas et al., 2016) was beneficial for families that were over or under represented in the sequencing data. Without the use of mFCFs, differences between the number of individuals caught in a haul (assessed morphologically) and the corresponding number of individuals calculated by back-estimation of RRA were as large as −8.92 (± SD 14.17) individuals on average for Ammodytidae, and 6.42 (± SD 7.78) individuals for Clupeidae. Once corrections had been applied, back-estimates of numbers of individuals of a given family reflected numbers of individuals assessed morphologically, differing, on average by less than 2 individuals per haul. No improvement was observed for families where the mean difference prior to applying an mFCF was less than 1 individual (Callionymidae and Triglidae, Table 1).

Correction factors are most useful for over or under represented taxa, if they are applicable regardless of the species composition or number of individuals within a bulk sample (Thomas et al., 2014, 2016). Here, each sample varied not only in species composition but also in total number of individuals (Table S1), suggesting that the application of correction factors can be beneficial for field studies, where input material varies in terms of abundance and composition. Further research is needed to see if mFCFs could be transferable from one study to another, however, using subsamples within a study to calculate family specific factors is likely to improve abundance estimates in the case of strongly over and underrepresented families.

Spatial patterns detected in community composition remained the same independent of whether they were assessed using morphological (CPUF) or back-estimated metabarcoding (RPUF). The small differences in abundance of rare taxa (Table S3), were mainly the result of miss-identification of *C. mustela* during morphological identification, indicating that metabarcoding of bulk samples may be used as a viable alternative to morphological identification of samples. All taxa detected in the survey are known to spawn in the survey area (Acevedo, Dwane, & Fives, 2002; Ellis et al., 2012). The family Ammodytidae is difficult to survey due to cryptic morphological characteristics (Ellis et al., 2012) and are therefore data limited. Here, we identified *A. marinus* and a species of the genus *Gymnammodytes*, further illustrating the potential for bulk metabarcoding for detecting cryptic species.

## Conclusions

We have shown that using a single marker (12S), equal amounts of tissue per sample and correction factors for amplification bias, and back-estimation of number of individuals, metabarcoding can provide accurate quantitative abundance estimates for the calculation of alpha and beta diversities. These methodological modifications could be applied to bulk samples from different terrestrial and marine habitats to improve abundance estimates. Specifically, we recommend the use of markers with highly conserved binding sites and using a small, equally sized pieces of tissue from each specimen to minimise biases and handling steps. Furthermore, where amplification bias does occur, the use of family level correction factors is recommended. This provides a rapid, community level assessment method, that could be used to further understand responses to disturbance and inter-annual or seasonal variability and monitor biodiversity in a changing global climate.

## Acknowledgements

We are very grateful to those who assisted with sampling: Helen MCCormick, Ross O’Neill, Michael Sheridan, Sarah Albuixech-Marti, Katie Costello and crew of the R.V. Celtic Voyager. This work has been funded by the European Regional Development fund through the Ireland Wales Co-operation Programme 2014 – 2020 (BlueFish project).

## Data Accessibility Statement

Sequences from metabarcoding have been deposited in the NCBI under accession reference BioProject PRJNA576002. Sanger sequences for the reference collection have been deposited in GenBank under accession numbers MN539918-MN539945 (CO-I) and MN539946-MN539976 (12S).

## Ethics approval

Sampling has been conducted following Home Office regulations and approved by Swansea University Ethics Committees under approval No. 181019/1996.

## Author contributions

SC, FR and CGL conceived the ideas. FR designed the methodology with help of DRB. FR and TUW carried out the laboratory work. FR performed bioinformatic analyses with TUW and RO. RO built the species database. FR led the writing of the manuscript with SC and all authors contributed critically to the drafts.

## References

Acevedo, S., Dwane, O., & Fives, J. M. (2002). The community structure of larval fish populations in an area of the Celtic Sea in 1998. Journal of the Marine Biological Association of the United Kingdom, 82(4), 641–648. doi:10.1017/S0025315402006008

Alberdi, A., Aizpurua, O., Gilbert, M. T. P., & Bohmann, K. (2018). Scrutinizing key steps for reliable metabarcoding of environmental samples. Methods in Ecology and Evolution, 9(1), 134–147. doi:10.1111/2041-210X.12849

Asch, R. G. (2015). Climate change and decadal shifts in the phenology of larval fishes in the California Current ecosystem. Proceedings of the National Academy of Sciences, 112(30), E4065–E4074. doi:10.1073/pnas.1421946112

Barange, M., Bahri, T., Beveridge, M.C.M., Cochrane, K.L., Funge-Smith, S. & Poulain, F., eds. 2018. Impacts of climate change on fisheries and aquaculture: synthesis of current knowledge, adaptation and mitigation options. FAO Fisheries and Aquaculture Technical Paper No. 627. Rome, FAO. 628 pp.

Becker, R. A., Sales, N. G., Santos, G. M., Santos, G. B., & Carvalho, D. C. (2015). DNA barcoding and morphological identification of neotropical ichthyoplankton from the Upper Paraná and São Francisco. Journal of Fish Biology, 87(1), 159–168. doi:10.1111/jfb.12707

Bohmann, K., Evans, A., Gilbert, M. T. P., Carvalho, G. R., Creer, S., Knapp, M., … de Bruyn, M. (2014). Environmental DNA for wildlife biology and biodiversity monitoring. Trends in Ecology and Evolution, 29(6), 358–367. doi:10.1016/j.tree.2014.04.003

Bolyen, E., Rideout, J. R., Dillon, M. R., Bokulich, N. A., Abnet, C. C., Al-Ghalith, G. A., … Caporaso, J. G. (2019). Reproducible, interactive, scalable and extensible microbiome data science using QIIME 2. Nature Biotechnology, 37(8), 852–857. doi:10.1038/s41587-019-0209-9

Borja, A., Elliott, M., Uyarra, M. C., Carstensen, J., & Mea, M. (2017). Editorial: Bridging the Gap between Policy and Science in Assessing the Health Status of Marine Ecosystems. Frontiers in Marine Science, 4(September). doi:10.3389/fmars.2017.00032

Boyer, F., Mercier, C., Bonin, A., Le Bras, Y., Taberlet, P. and Coissac, E., 2015. obitools: a unix-inspired software package for DNA metabarcoding. Molecular Ecology Resources, 1(16), pp. 176–182. doi:10.1111/1755-0998.12428

Brechon, A. L., Coombs, S. H., Sims, D. W., & Griffiths, A. M. (2013). Development of a rapid genetic technique for the identification of clupeid larvae in the Western English Channel and investigation of mislabelling in processed fish products. ICES Journal of Marine Science, 70(2), 399–407. doi:10.1093/icesjms/fss178

Callahan, B. J., McMurdie, P. J., & Holmes, S. P. (2017). Exact sequence variants should replace operational taxonomic units in marker-gene data analysis. ISME Journal, 11(12), 2639–2643. doi:10.1038/ismej.2017.119

Canfield, T. J., & Jones, J. R. (1996). Zooplankton abundance, biomass, and size-distribution in selected midwestern waterbodies and relation with trophic state. Journal of Freshwater Ecology, 11(2), 171–181. doi:10.1080/02705060.1996.9663476

Clarke, K. R. (1993). Non-parametric multivariate analyses of changes in community structure. Australian Journal of Ecology, 18(1), 117–143. doi:10.1111/j.1442-9993.1993.tb00438.x

Collins, R. A., Bakker, J., Wangensteen, O. S., Soto, A. Z., Corrigan, L., Sims, D. W., … Mariani, S. (2019). Non-specific amplification compromises environmental DNA metabarcoding with COI. Methods in Ecology and Evolution, 2041–210X.13276. doi:10.1111/2041-210X.13276

Deagle, B. E., Jarman, S. N., Coissac, E., Pompanon, F., & Taberlet, P. (2014). DNA metabarcoding and the cytochrome c oxidase subunit I marker: not a perfect match DNA metabarcoding and the cytochrome c oxidase subunit I marker: not a perfect match. Biology Letters, 10(September), 2–5. doi:10.1098/rsbl.2014.0562

Deagle, B. E., Thomas, A. C., McInnes, J. C., Clarke, L. J., Vesterinen, E. J., Clare, E. L., … Eveson, J. P. (2019). Counting with DNA in metabarcoding studies: How should we convert sequence reads to dietary data? Molecular Ecology, 28(2), 391–406. doi:10.1111/mec.14734

Edgar RC. 2010. Search and clustering orders of magnitude faster than BLAST. Bioinformatics 26:2460–2461. DOI: 10.1093/bioinformatics/btq461.

Elbrecht, V., Peinert, B., & Leese, F. (2017). Sorting things out: Assessing effects of unequal specimen biomass on DNA metabarcoding. Ecology and Evolution, 7(17), 6918–6926. doi:10.1002/ece3.3192

Ellis, J. R., Milligan, S. P., Readdy, L., Taylor, N., & Brown, M. J. (2012). Spawning and nursery grounds of selected fish species in UK waters. Biological Conservation, (147), 60. Retrieved from https://www.cefas.co.uk/publications/techrep/TechRep147.pdf

Findley, K., Oh, J., Yang, J., Conlan, S., Deming, C., Meyer, J. A., … Segre, J. A. (2013). Topographic diversity of fungal and bacterial communities in human skin. Nature, 498(7454), 367–370. doi:10.1038/nature12171

Gillet, B., Cottet, M., Destanque, T., Kue, K., Descloux, S., Chanudet, V., & Hughes, S. (2018). Direct fishing and eDNA metabarcoding for biomonitoring during a 3-year survey significantly improves number of fish detected around a South East Asian reservoir. PLoS ONE, 13(12), 1–25. doi:10.1371/journal.pone.0208592

Hernandez, F. J., Carassou, L., Graham, W. M., & Powers, S. P. (2013). Evaluation of the taxonomic sufficiency approach for ichthyoplankton community analysis. Marine Ecology Progress Series, 491(Fahay 2007), 77–90. doi:10.3354/meps10475

Hunt, B. P. V, & Hosie, G. W. (2003). The Continuous Plankton Recorder in the Southern Ocean: A comparative analysis of zooplankton communities sampled by the CPR and vertical net hauls along 140°E. Journal of Plankton Research, 25(12), 1561–1579. doi: 10.1093/plankt/fbg108

Huson, D. H., Auch, A. F., Qi, J., & Schuster, S. C. (2007). MEGAN analysis of metagenomic data. Genome Research, 17(3), 377–386. doi:10.1101/gr.5969107

Kimmerling, N., Zuqert, O., Amitai, G., Gurevich, T., Armoza-Zvuloni, R., Kolesnikov, I., … Sorek, R. (2018). Quantitative species-level ecology of reef fish larvae via metabarcoding. Nature Ecology & Evolution, 2(2), 306–316. doi:10.1038/s41559-017-0413-2

Krehenwinkel, H., Wolf, M., Lim, J. Y., Rominger, A. J., Simison, W. B., & Gillespie, R. G. (2017). Estimating and mitigating amplification bias in qualitative and quantitative arthropod metabarcoding. Scientific Reports, 7(1). doi:10.1038/s41598-017-17333-x

Lamb, P. D., Hunter, E., Pinnegar, J. K., Creer, S., Davies, R. G., & Taylor, M. I. (2019). How quantitative is metabarcoding: A meta-analytical approach. Molecular Ecology, 28(2), 420–430. doi:10.1111/mec.14920

Morgulis, A., Coulouris, G., Raytselis, Y., Madden, T. L., Agarwala, R., & Schäffer, A. A. (2008). Database indexing for production MegaBLAST searches. Bioinformatics, 24(16), 1757–1764. doi:10.1093/bioinformatics/btn322

Nobile, A. B., Freitas-Souza, D., Ruiz-Ruano, F. J., Nobile, M. L. M. O., Costa, G. O., de Lima, F. P., … Oliveira, C. (2019). DNA metabarcoding of Neotropical ichthyoplankton: Enabling high accuracy with lower cost. Metabarcoding and Metagenomics, 3, 69–76. doi:10.3897/mbmg.3.35060

Piñol, J., Mir, G., Gomez-Polo, P., & Agustí, N. (2015). Universal and blocking primer mismatches limit the use of high-throughput DNA sequencing for the quantitative metabarcoding of arthropods. Molecular Ecology Resources, 15(4), 819–830. doi:10.1111/1755-0998.12355

Piñol, J., Senar, M. A., & Symondson, W. O. C. (2019). The choice of universal primers and the characteristics of the species mixture determine when DNA metabarcoding can be quantitative. Molecular Ecology, 28(2), 407–419. doi:10.1111/mec.14776

Planque, B., & Frédou, T. (1999). Temperature and the recruitment of Atlantic cod (Gadus morhua). Canadian Journal of Fisheries and Aquatic Sciences, 56(11), 2069–2077. doi:10.1139/f99-114

Porter, T. M., & Hajibabaei, M. (2018). Scaling up: A guide to high-throughput genomic approaches for biodiversity analysis. Molecular Ecology, 27(2), 313–338. doi:10.1111/mec.14478

Radchuk, V., Turlure, C., & Schtickzelle, N. (2013). Each life stage matters: The importance of assessing the response to climate change over the complete life cycle in butterflies. Journal of Animal Ecology, 82(1), 275–285. doi:10.1111/j.1365-2656.2012.02029.x

[dataset] Ratcliffe F. C., Ratcliffe, Uren Webster, T. M., Rodriguez-Barreto, D., O’Rorke, R., Garcia de Leaniz, C., Consuegra, S. (2019); Rapid methods for quantitative fish larvae community assessment using metabarcoding; NCBI SRA; BioProject: PRJNA576002 and GenBank accession numbers MN539918-MN539945 (CO-I) and MN539946–MN539976 (12S).

Riaz, T., Shehzad, W., Viari, A., Pompanon, F., Taberlet, P., & Coissac, E. (2011). EcoPrimers: Inference of new DNA barcode markers from whole genome sequence analysis. Nucleic Acids Research, 39(21), 1–11. doi:10.1093/nar/gkr732

Richardson, A. J., Walne, A. W., John, A. W. G., Jonas, T. D., Lindley, J. A., Sims, D. W., … Witt, M. (2006). Using continuous plankton recorder data. Progress in Oceanography, 68(1), 27–74. doi:10.1016/j.pocean.2005.09.011

Richardson, R. T., Lin, C., Quijia, J. O., Riusech, N. S., Goodell, K., & Johnson, R. M. (2015). Rank-based characterization of pollen assemblages collected by honey bees using a multi-locus metabarcoding approach. Applications in Plant Sciences, 3(11), 1500043. doi: 10.3732/apps.1500043

Rodriguez, J. M., Alemany, F., & Garcia, A. (2017). A guide to the eggs and larvae of 100 common Western Mediterranean Sea bony fish species.

Russell, F. S. (1976). The eggs and planktonic stages of British marine fishes (Vol. 524). London: Academic press.

Schloss, P. D., Westcott, S. L., Ryabin, T., Hall, J. R., Hartmann, M., Hollister, E. B., … Weber, C. F. (2009). Introducing mothur: Open-source, platform-independent, community-supported software for describing and comparing microbial communities. Applied and Environmental Microbiology, 75(23), 7537–7541. doi:10.1128/AEM.01541-09

Schnell, I. B., Bohmann, K., & Gilbert, M. T. P. (2015). Tag jumps illuminated - reducing sequence-to-sample misidentifications in metabarcoding studies. Molecular Ecology Resources, 15(6), 1289–1303. doi:10.1111/1755-0998.12402

Sigut, M., Kostovćik, M., Sigutova, H., Hulcr, J., Drozd, P., & Hrcek, J. (2017). Performance of DNA metabarcoding, standard barcoding, and morphological approach in the identification of hostparasitoid interactions. PLoS ONE, 12(12), 1–18. doi:10.1371/journal.pone.0187803

Taberlet, P., Coissac, E., Pompanon, F., Brochmann, C., & Willerslev, E. (2012). Towards next-generation biodiversity assessment using DNA metabarcoding. Molecular Ecology, 21(8), 2045–2050. doi:10.1111/j.1365-294X.2012.05470.x

Tang, M., Hardman, C. J., Ji, Y., Meng, G., Liu, S., Tan, M., … Yu, D. W. (2015). High-throughput monitoring of wild bee diversity and abundance via mitogenomics. Methods in Ecology and Evolution, 6(9), 1034–1043. doi:10.1111/2041-210X.12416

Thomas, A. C., Deagle, B. E., Eveson, J. P., Harsch, C. H., & Trites, A. W. (2016). Quantitative DNA metabarcoding: Improved estimates of species proportional biomass using correction factors derived from control material. Molecular Ecology Resources, 16(3), 714–726. doi:10.1111/1755-0998.12490

Thomas, A. C., Jarman, S. N., Haman, K. H., Trites, A. W., & Deagle, B. E. (2014). Improving accuracy of DNA diet estimates using food tissue control materials and an evaluation of proxies for digestion bias. Molecular Ecology, 23(15), 3706–3718. doi:10.1111/mec.12523

Thomsen, P. F., Møller, P. R., Sigsgaard, E. E., Knudsen, S. W., Jørgensen, O. A., & Willerslev, E. (2016). Environmental DNA from seawater samples correlate with trawl catches of subarctic, deepwater fishes. PLoS ONE, 11(11). doi:10.1371/journal.pone.0165252

Ward, R. D., Zemlak, T. S., Innes, B. H., Last, P. R., & Hebert, P. D. N. (2005). DNA barcoding Australia’s fish species. Philosophical Transactions of the Royal Society B: Biological Sciences, 360(1462), 1847–1857. doi:10.1098/rstb.2005.1716

