## Supplementary material for "Rapid quantitative assessment of fish larvae community composition using metabarcoding"

Table S1. Number of individual larvae captured in each haul in the survey.

| Haul number | Total individuals in haul |
|---|---|
| 1 | 1 |
| 2 | 63 |
| 3 | 59 |
| 4 | 52 |
| 5 | 27 |
| 6 | 1 |
| 7 | 58 |
| 8 | 32 |
| 9 | 8 |
| 10 | 0 |
| 11 | 0 |
| 12 | 24 |
| 13 | 6 |
| 14 | 0 |

Table S2. Overview of ichthyoplankton taxonomic assignment. Detection (presence/absence): families assigned by morphology alone, morphology corrected by a subsample of Sanger sequencing, and 12S metabarcoding. ‘x’ indicates where a method achieved lowest taxonomic classification. NA indicates where a taxon was unidentified by a method. Abundance: the total number of individuals detected by sanger corrected morphology and total number of bulk reads post filtering.

| Taxonomic classification | | Detection | | | Abundance | |
|---|---|---|---|---|---|---|
| Family | Lowest classification | Morphology alone | Sanger Corrected morphology | bulk | Sanger Corrected morphology | No. reads (post filtering) |
| f__Ammodytidae | f__Ammodytidae | Ammodytidae | x | x | 5 | 24429 |
| | s__Ammodytes_marinus | | x | x | 26 | 247248 |
| | g__Gymnammodytes | | | x | | 90543 |
| f__Callionymidae | g__Callionymus | Callionymidae | x | x | 62 | 586077 |
| f__Clupeidae | x__C. harengus_ S.sprattus | Cluepeidae | x | Clupea/Sprattus | 128 | 427286 |
| | s__Sardina_pilchardus | *NA* | *NA* | x | 0 | 8784 |
| f__Gadidae | s__Merlangius_merlangus | x | x | Pollaccius/Merlangius spp | 9 | 0 |
| | s__Micromesistius_poutassou | x | x | *NA* | 5 | 0 |
| | x__P.pollachius/virens_M. merlangus | *NA* | *NA* | Pollaccius/Merlangius spp | 0 | 131238 |
| | s__Pollachius_pollachius | x | x | Pollaccius/Merlangius spp | 2 | 0 |
| | s__Pollachius_virens | x | x | Pollaccius/Merlangius spp | 1 | 0 |
| | g__Tricopterus | x | *NA* | *NA* | 0 | 0 |
| | s__Trisopterus_esmarkii | x | x | x | 6 | 93440 |
| | s__Trisopterus_minutus | x | x | x | 21 | 122247 |
| f__Gaidropsaridae | s__Ciliata_mustela | *NA* | x | x | 5 | 183976 |

| | | | | | | |
|---|---|---|---|---|---|---|
| f__Gobiidae | f__Gobiidae | Gobiidae | *NA* | x | 0 | 3168 |
| | s__Buenia_jeffreysii | *NA* | x | x | 1 | 8306 |
| | s__Aphia_minuta | Aphia minuta | *NA* | *NA* | 0 | 0 |
| f__Labridae | s__Labrus_bergylta | *NA* | x | x | 3 | 20578 |
| | s__Labrus_mixtus | *NA* | *NA* | x | 0 | 5172 |
| f__Lotidae | s__Molva_molva | *NA* | *NA* | x | 0 | 14595 |
| f__Merlucciidae | s__Merluccius_merluccius | x | x | x | 2 | 104615 |
| f__Mugilidae | f__Mugilidae | x | *NA* | *NA* | 0 | 0 |
| f__Pleuronectidae | f__Pleuronectidae | x | x | x | 0 | 117330 |
| | s__Glyptocephalus_cynoglossus | x | x | Pleuronectidae | 12 | 0 |
| | s__Limanda_limanda | x | x | x | 13 | 129542 |
| | s__Microstomus_kitt | x | x | x | 11 | 142793 |
| f__Scophthalmidae | g__Lepidorhombus_sp | *NA* | x | x | 3 | 26632 |
| f__Solidae | f__Solidae | x | *NA* | *NA* | 0 | 0 |
| f__Triglidae | f__Triglidae | *NA* | x | x | 9 | 28714 |
| damaged | damaged | *NA* | x | 0 | 8 | 0 |
| unknown | unknown | x | *NA* | *NA* | 0 | 0 |

Table S3. SIMPER analysis showing average abundances (Av.Abund), average dissimilarity between locations (Av.Diss), the contribution dissimilarity between locations (Contrib%) and the cumulative contributions dissimilarity (Cum.%) for each of the 7 taxa which contribute the most to between group dissimilarity between locations 2 and 3, using morphology and metabarcoding.

| Species | Location 2 Av.Abund | Location 3 Av.Abund | Av.Diss | Contrib% | Cum.% |
|---|---|---|---|---|---|
| **Morphology** | | | | | |
| x__C. harengus_S. sprattus | 0.23 | 0.02 | 18.77 | 21.44 | 21.44 |
| f__Triglidae | 0.05 | 0.02 | 12.58 | 14.37 | 35.8 |
| s__Callionymus_ukspp | 0.11 | 0.11 | 10.99 | 12.54 | 48.35 |
| s__Microstomus_kitt | 0.06 | 0.05 | 5.92 | 6.76 | 55.11 |
| s__Trisopterus_minutus | 0.05 | 0.03 | 5.46 | 6.23 | 61.33 |
| s__Ciliata_mustela | 0.06 | 0 | 4.82 | 5.5 | 66.84 |
| S__Merlangius _merlangus | 0.02 | 0.06 | 4.63 | 5.29 | 72.12 |
| **Back-estimated reads** | | | | | |
| x__C. harengus_S. sprattus | 0.21 | 0.03 | 17.28 | 19.91 | 19.91 |
| f__Triglidae | 0.04 | 0.02 | 12.19 | 14.05 | 33.96 |
| s__Callionymus_ukspp | 0.09 | 0.09 | 9.47 | 10.91 | 44.87 |
| s__Ciliata_mustela | 0.09 | 0 | 6.95 | 8.01 | 52.88 |
| s__Microstomus_kitt | 0.05 | 0.05 | 5.68 | 6.54 | 59.42 |
| s__Trisopterus_minutus | 0.06 | 0.03 | 5.35 | 6.16 | 65.58 |
| s__Limanda_limanda | 0.04 | 0.02 | 5.27 | 6.07 | 71.64 |

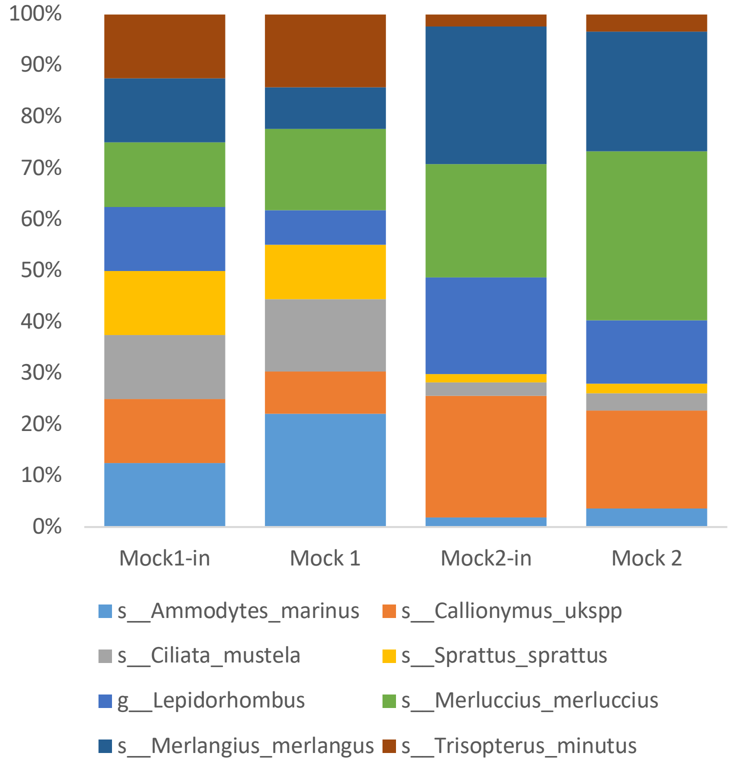


Figure S1. Metabarcoding relative read abundance in two mock communities (Mock 1, Mock 2) constructed of equimolar quantities of Sanger-barcoded genomic DNA (Mock1-in), and varying molarities of genomic DNA (Mock2-in).

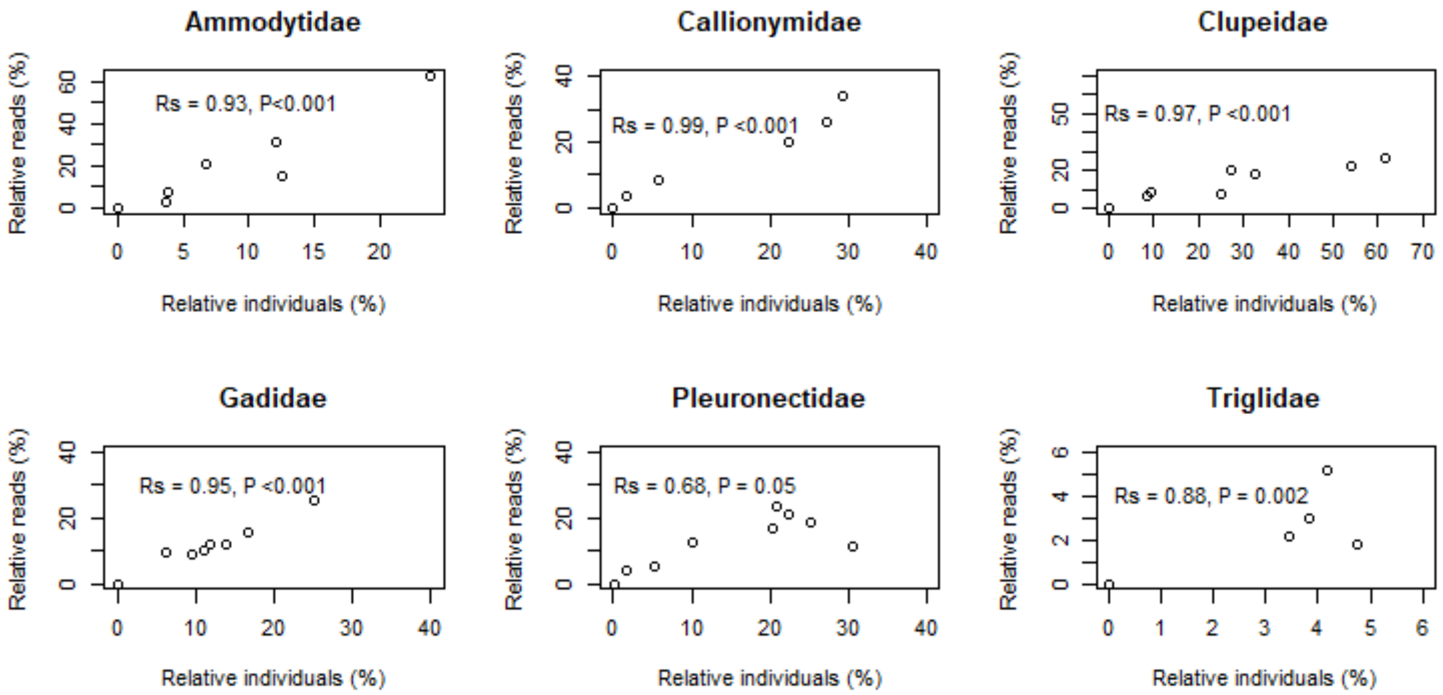


Figure S2. Relationship between relative number of individuals (%) within a taxonomic family in each haul, and relative number of reads post filtering (%) in the corresponding sample. Family level Spearman's Rho correlation were calculated across all hauls in the survey.

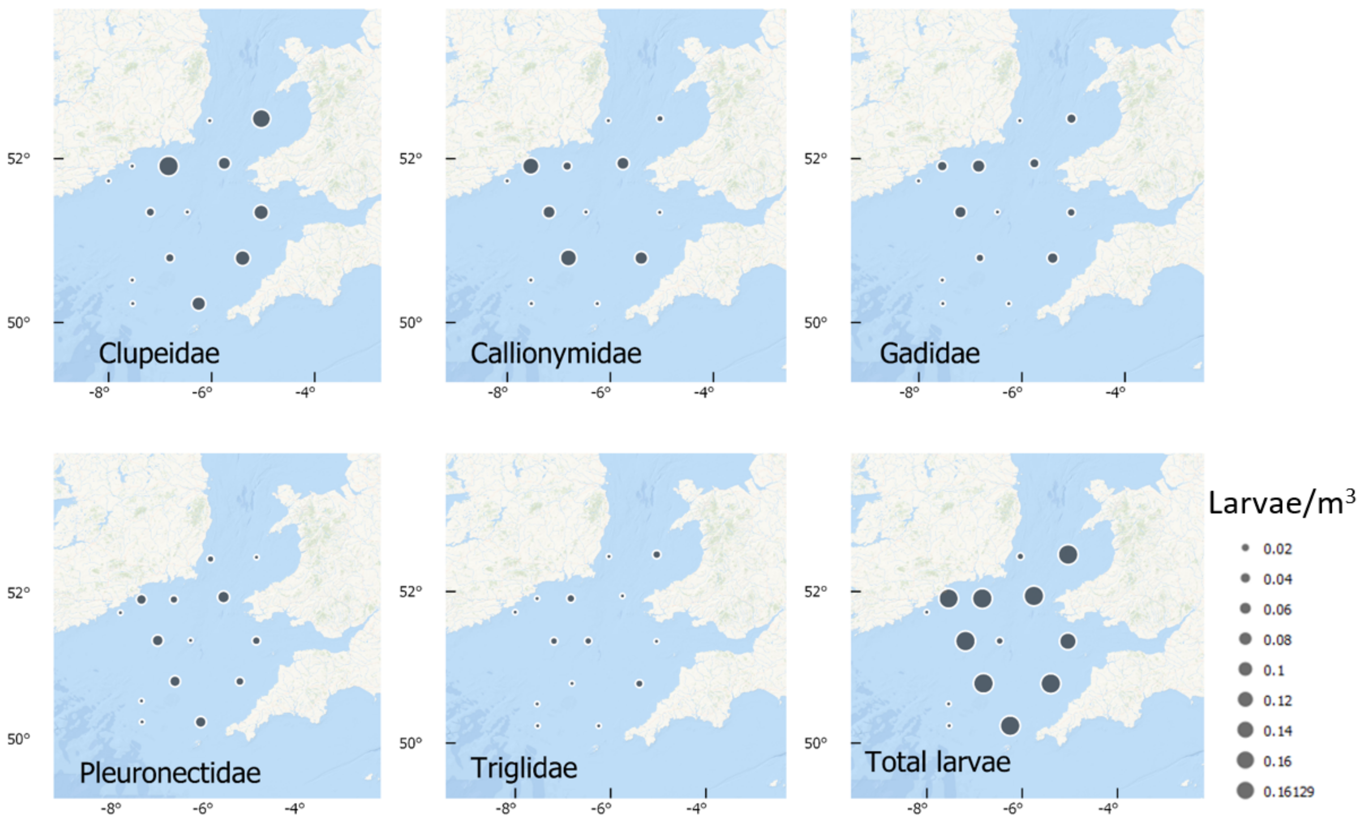


Figure S3. Abundances (number individuals) of families per $M^3$ of water filtered within survey, assessed by morphology.

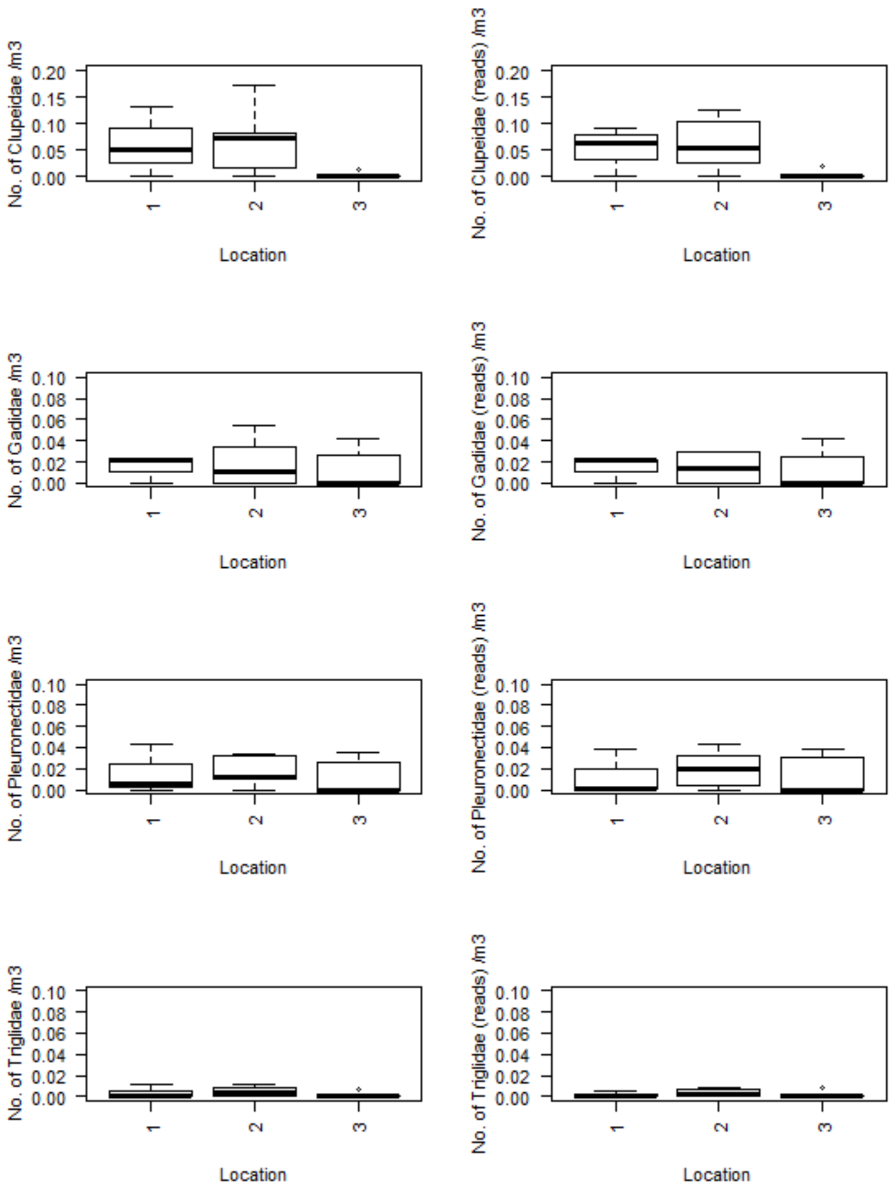


Figure S4. Abundance of individuals of a given family per m$^3$ in each of the 3 locations, based on morphology (left, CPUF) and back-estimated reads (right, RPUF).
